# Cell-type specific auditory responses in the striatum are shaped by feed forward inhibition

**DOI:** 10.1101/2024.05.09.592848

**Authors:** Mélanie Druart, Megha Kori, Corryn Chaimowitz, Catherine Fan, Tanya Sippy

## Abstract

The posterior “tail” region of the striatum receives dense innervation from sensory brain regions and has been demonstrated to play a role in behaviors that require sensorimotor integration including discrimination^1,2^, avoidance^3^ and defense^4^ responses. The output neurons of the striatum, the D1 and D2 striatal projection neurons (SPNs) that make up the direct and indirect pathways, respectively, are thought to play differential roles in these behavioral responses, although it remains unclear if or how these neurons display differential responsivity to sensory stimuli. Here, we used whole-cell recordings in vivo and ex vivo to examine the strength of excitatory and inhibitory synaptic inputs onto D1 and D2 SPNs following the stimulation of upstream auditory pathways. While D1 and D2 SPNs both displayed stimulus-evoked depolarizations, D1 SPN responses were stronger and faster for all stimuli tested in vivo as well as in brain slices. This difference did not arise from differences in the strength of excitatory inputs but from differences in the strength of feed forward inhibition. Indeed, fast spiking interneurons, which are readily engaged by auditory afferents exerted stronger inhibition onto D2 SPNs compared to D1 SPNs. Our results support a model in which differences in feed forward inhibition enable the preferential recruitment of the direct pathway in response to auditory stimuli, positioning this pathway to initiate sound-driven actions.

## Introduction

The ability to use sensory information to guide our actions is a fundamental ability of brain circuitry that is crucial for survival. The input nucleus of the basal ganglia, the striatum, is thought to integrate contextual, motor and reward information to control action initiation and selection. Several anatomical investigations have confirmed the presence and organization of functional sensory inputs throughout the rodent dorsal striatum^1–3^. It is well established, for example, that primary sensory cortical areas and sensory thalamic nuclei project to this region, making glutamatergic connections onto striatal neurons in a cell-type specific manner^4^. However, it remains unclear how this connectivity is related to the responsiveness of these neurons to sensory stimuli remains unclear.

Striatal neurons are predominantly GABAergic striatal projection neurons (SPNs) that can be divided into two main classes depending on their projection patterns and expression of dopamine receptors: D1-expressing SPNs, (D1 SPNs) make up the “direct” pathway, and directly project to the output nuclei of the basal ganglia, while D2-expressing SPNs (D2 SPNs) make up the “indirect” pathway, and project to globus pallidus which then projects to the output nuclei. D1 SPN activity has been classically associated with increased movement, while D2 SPN activity is associated with reduced movement^5^. Yet, pathway-specific differences in the activity of these neural populations have been hard to reveal during spontaneous movements and/or locomotion^6–9^. In contrast, we previously reported that during sensory-evoked movements, D1 SPNs are preferentially activated before D2 SPNs and that this earlier activation might serve to initiate sensory-triggered actions^10^. Here, we investigate the synaptic responses of striatal neurons resulting from sensory inputs both in vivo and ex vivo. We focused on the posterior striatum (pStr) that gets rich sensory input, particularly from auditory areas such as primary auditory cortex (A1) and auditory thalamus (MGB).

The pStr has been shown to be important for auditory decision making, with inhibition of the region^11^, or its dopaminergic inputs^12^ leading to dampened choice performance. Notably, silencing of dopamine neurons was shown to inhibit SPN responses to auditory stimuli while only selective blockade of D1 receptors, not D2 receptors, dampened performance^12^. In other studies, dopaminergic inputs to the pStr have been implicated in mediating behavioral responses to threatening stimuli ^13–16^. In these studies, activity of D1 SPNs is more tightly correlated with the presentation of a threatening stimulus, and is important for learning, whereas D2 SPN activity mediates freezing and retreat behavior in the absence of a learned conditioned stimulus^15^. However, the sensory-evoked response properties onto these cell types have not been thoroughly investigated, either ex vivo or in vivo, leaving it unclear how the two pathways process sensory information to mediate behavior. Specifically, it is unclear if the D1 and D2 pathways are differentially sensitive to sensory inputs.

To investigate this, we combined whole-cell recordings with optogenetics to distinguish D1 from D2 SPNs (i.e., “optopatch” technique)^17^ and measured their membrane potential (V_m_) to auditory stimuli in awake and anesthetized mice. We report that auditory-evoked responses were more robust in D1 SPNs compared with D2 SPNs. Ex vivo experiments revealed that, although D1 and D2 SPNs receive comparably strong excitatory inputs from auditory afferents, feed forward inhibition is stronger onto D2 SPNs. Indeed, we found that local parvalbumin (PV) interneurons make stronger connections onto D2 SPNs compared to D1 SPNs. Our results show that in the pStr, direct pathway neurons respond more robustly to auditory stimuli, a difference that is supported by differences in the local network connectivity of fast spiking interneurons onto direct and indirect pathway neurons.

## Results

### In vivo auditory-evoked synaptic responses in D1 and D2 SPNs

We played auditory stimuli (7 chords, with center frequencies ranging from 5.1 to 31.9Hz, logarithmically spaced) to head-fixed awake and anesthetized mice and simultaneously obtained whole-cell V_m_ recordings in the pStr (Figure 1A). The whole-cell recording technique allows for the measurement of subthreshold synaptic potentials and enables the quantification of cumulative connection strength of afferent areas that are recruited by auditory stimuli^18^. Post-hoc biocytin labeling allowed us to determine the precise location of recorded neurons in the pStr (Figures 1B, S1A, D). We employed mice that expressed Channelrhodopsin-2 (ChR2) in D2 neurons (D2-Cre^19^ x Ai32^20^) and used the “optopatch” method to distinguish ChR2-positive D2 SPNs from ChR2-negative putative D1 SPNs^17,21,22^. In brief, after a stable recording was established, 500 ms long pulses of blue light were applied via the patch pipette, resulting in short-latency depolarizations and action potential (AP) firing in neurons expressing ChR2 (Figure 1C). D2 SPNs were therefore all light-responsive and could be differentiated from other interneuron types by their low firing rates and hyperpolarized resting V_m_^23^. ChR2-negative neurons showed mostly no response to light or had a slight hyperpolarization due to recurrent connectivity from nearby ChR2-positive D2 SPNs, which are GABAergic. They also had properties consistent with SPNs, such as low firing rates and high input resistances making them distinguishable from the minority of other cell types in this region^23,24^.

**Figure 1:**
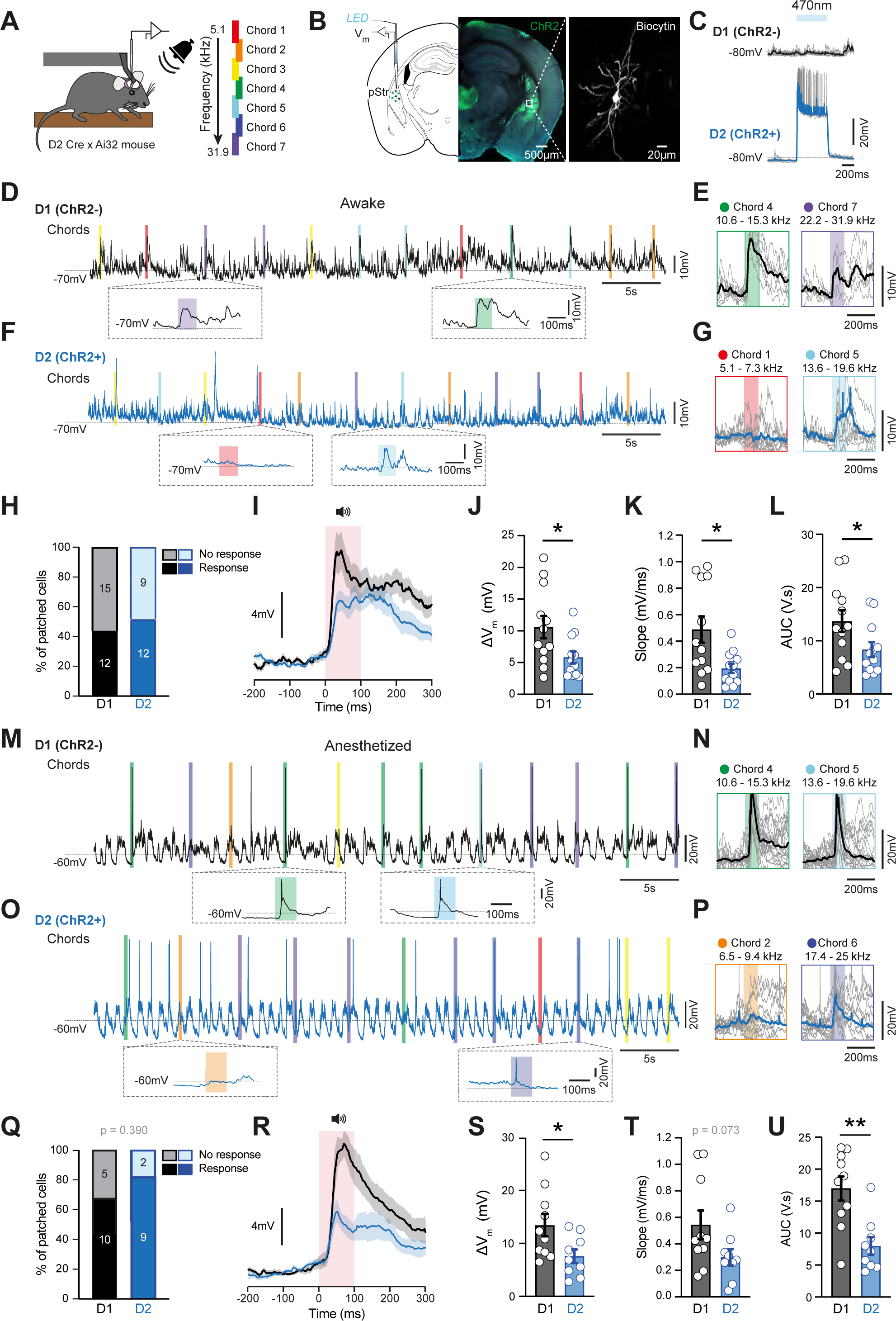
Cell-Type Specific Auditory-evoked Responses in the pStr. (A) Mice were head-restrained for V_m_ recordings during sound presentation of 7 discreet chords with center frequencies ranging from 5.1 to 31.9 kHz that were played at random time intervals (3-5 s). (B) Neurons were targeted for recordings in the pStr of D2-ChR2-YFP-mice (left), which enabled optogenetic identification using the optopatcher. Patched neurons were filled with biocytin to allow for post-hoc identification of the cell and its precise location (right). (C) Examples of responses of neurons to 500 ms light stimulation delivered through the optopatcher. Top: A ChR2-negative putative D1 SPN cell that did not respond with depolarization in response to a 500 ms blue light stimulation. Bottom: a ChR2-positive D2 SPN that did respond with depolarization. (D) Example trace from a D1 SPN showing subthreshold V_m_ activity. Vertical colored bars indicate chord presentations. (E) V_m_ of the D1 SPN shown in D for all trials in response to two chords (black average, grey individual; chord 4: n = 8 trials, chord 7: n = 6 trials). (F) Example trace from a D2 SPN. (G) V_m_ of the D2 SPN in F (blue average, gray individual; chord 1: n = 10 trials, chord 5: n = 7 trials) (H) Percent of D1 and D2 SPNs responsive to at least one chord (p = 0.56) (I) Grand average V_m_ across all responsive D1 (black, n = 12) and D2 (blue, n = 12) SPNs of the chord in each neuron that yielded the maximum response. (J) The maximum delta V_m_ (calculated as the peak depolarization during the first 50 ms after stimulus presentation) was larger in D1 SPNs (p = 0.026). (K) The slope of the stimulus-triggered response was significantly larger in D1 SPNs than D2 SPNs (p = 0.011). (L) The area under the curve (AUC) during the first 50 ms was larger in D1 SPNs than D2 SPNs (p = 0.040). (M) Example of a V_m_ trace in a D1 SPN in anesthetized mice, vertical bars represent chord presentations. (N) V_m_ of the D1 SPN shown in M for all trials in response to two chords (black average, grey individual, chord 4: n = 20 trials, chord 5: n = 18 trials) (O) Same as M, for a D2 SPN. (P) Same as N, for a D2 SPN (blue average, grey individual, chord 2: n = 12 trials, chord 6: n = 14 trials). (Q) percent of D1 and D2 SPNs responsive to at least 1 chord under anesthesia (p = 0.390) (R) Grand average V_m_ across all responsive D1 (black, n = 10) and D2 (blue, n = 9) SPNs of the chord in each neuron that yielded the maximum response. (S) The maximum delta V_m_ (calculated as the peak depolarization during the first 50 ms after stimulus presentation) was larger in D1 SPNs in anesthetized mice (p = 0.032). (T) The slope of the stimulus-triggered response was not significantly larger between D1 and D2 SPNs (p = 0.073). (U) The AUC of the stimulus-triggered response was significantly larger in D1 SPNs (p = 0.003). All data are represented as mean ± SEM. Each open circle represents an individual cell. * p < 0.05, ** p < 0.01.

We recorded from 21 D2 SPNs and 27 D1 SPNs in awake mice, all of which displayed spontaneous subthreshold V_m_ dynamics (Figure 1D-G). Of these, 12/21 (57 %) D2 SPNs and 12/27 (44 %) D1 SPNs were characterized as having a response to at least 1 chord, with no difference in these proportions (Figure 1H, p = 0.561, Chi-square test). In this subset, we found that V_m_ responses within 0-60 ms of the stimulus were larger in D1 SPNs compared to D2 SPNs (Figure 1I). Indeed, V_m_ deflections in D1 SPNs were on average larger (Figure 1J; maximum delta V_m_: D1 = 10.58 ± 1.76 mV; D2 = 5.84 ± 0.95 mV, p = 0.026; Student’s t-test) and faster (Figure 1K; rate of rise, or slope: D1 = 0.489 ± 0.099 mV/ms; D2 = 0.200 ± 0.124 mV/ms, p = 0.011; Student’s t-test), cumulating into a greater overall depolarization (Figure 1L; area under the curve (AUC): D1 = 13.7 ± 2.02 V.s; D2 = 8.36 ± 1.39 V.s, p = 0.040, Student’s t-test). D1 SPNs also responded to significantly more chords than D2 SPNs (Figure S1B-C; cell-type factor: 0.0182; 1^st^ p = 0.0209; 2^nd^ p = 0.041; 3^rd^ p = 0.0351; 4^th^ p = 0.089; Two-way RM ANOVA).

Given that sensory-evoked movements may contribute to the observed responses of striatal neurons^22^, we repeated these experiments in mice anesthetized with ketamine/xylazine. These recordings revealed UP and DOWN states of the V_m_ that are typical of anesthesia^17,21,25–27^. Auditory responses could be evoked in both states (Figure 1M-P). Similar to our findings in awake mice, the proportion of auditory-responsive cells did not differ between D1 and D2 SPNs, with 10/15 D1 SPNs and 9/11 D2 SPNs showing responses (Figure 1Q, p = 0.390; Chi-square test). In addition, the average V_m_ response of D1 SPNs was larger than D2 SPNs (Figure 1R), and demonstrated a larger maximum delta V_m_ (Figure 1S; D1 = 13.47 ± 2.08 mV; D2 = 7.60 ± 1.28 mV, p = 0.032; Student’s t-test), a faster rate of rise (Figure 1T; D1 = 0.544 ± 0.108 mV/ms; D2 = 0.299 ± 0.061 mV/ms, p = 0.073; Student’s t-test) and larger overall response, as measured by the AUC (Figure 1U; D1 = 16.99 ± 1.90 V.s; D2 = 7.99 ± 1.38 V.s, p = 0.003; Mann Whitney). As in awake animals, D1 SPNs tended to respond to more chords than D2 SPNs (Figure S1F). Therefore, even in the absence of movement, D1 SPNs demonstrated larger auditory-evoked responses than D2 SPNs.

### Cell-type specific connectivity from auditory thalamus and cortex onto D1 and D2 SPNs

The cell-type specific responses observed in vivo may be attributable to differences in intrinsic excitability between direct and indirect pathway neurons. However, similar to previous reports, we found that D2 SPNs are more excitable than D1 SPNs in vivo (Figure S1G-H), a difference that would render them being more rather than less responsive to inputs. It could also arise from differences in connectivity from upstream auditory brain regions such as the medial geniculate nucleus (MGB) of the thalamus and A1 of the cortex. To test this, we made use of rabies viruses to transsynaptically trace afferent inputs to both SPN cell types^28^. We injected helper viruses (AAV-FLEX-TVA and AAV-FLEX-N2cG) in the pStr of either D1-Cre or A2A-Cre mice. We later injected pseudotyped, G-deficient CVS-N2c rabies expressing GFP^29^ (EnVA-N2cΔG-GFP) in the same region (Figure 2A. S2A). This resulted in intense GFP expression in both D1-Cre and A2A-Cre mice in MGB and auditory cortex (AudCx, including primary and secondary auditory cortices) (Figures 2B, S2B). We then calculated the percent of neurons from AudCx and MGB projecting to D1 or D2 SPNs, finding no difference from either region (Figures 2C, S2B-D; AudCx: D1-Cre = 10.83 ± 1.31, n = 4 mice, A2A-Cre = 15.34 ± 1.88, n = 4 mice; MGB : D1-Cre = 12.13 ± 0.68, n = 4 mice A2A-Cre = 11.09 ± 0.64, n = 4 mice; cell-type factor, p = 0.19, Two-Way RM ANOVA).

**Figure 2:**
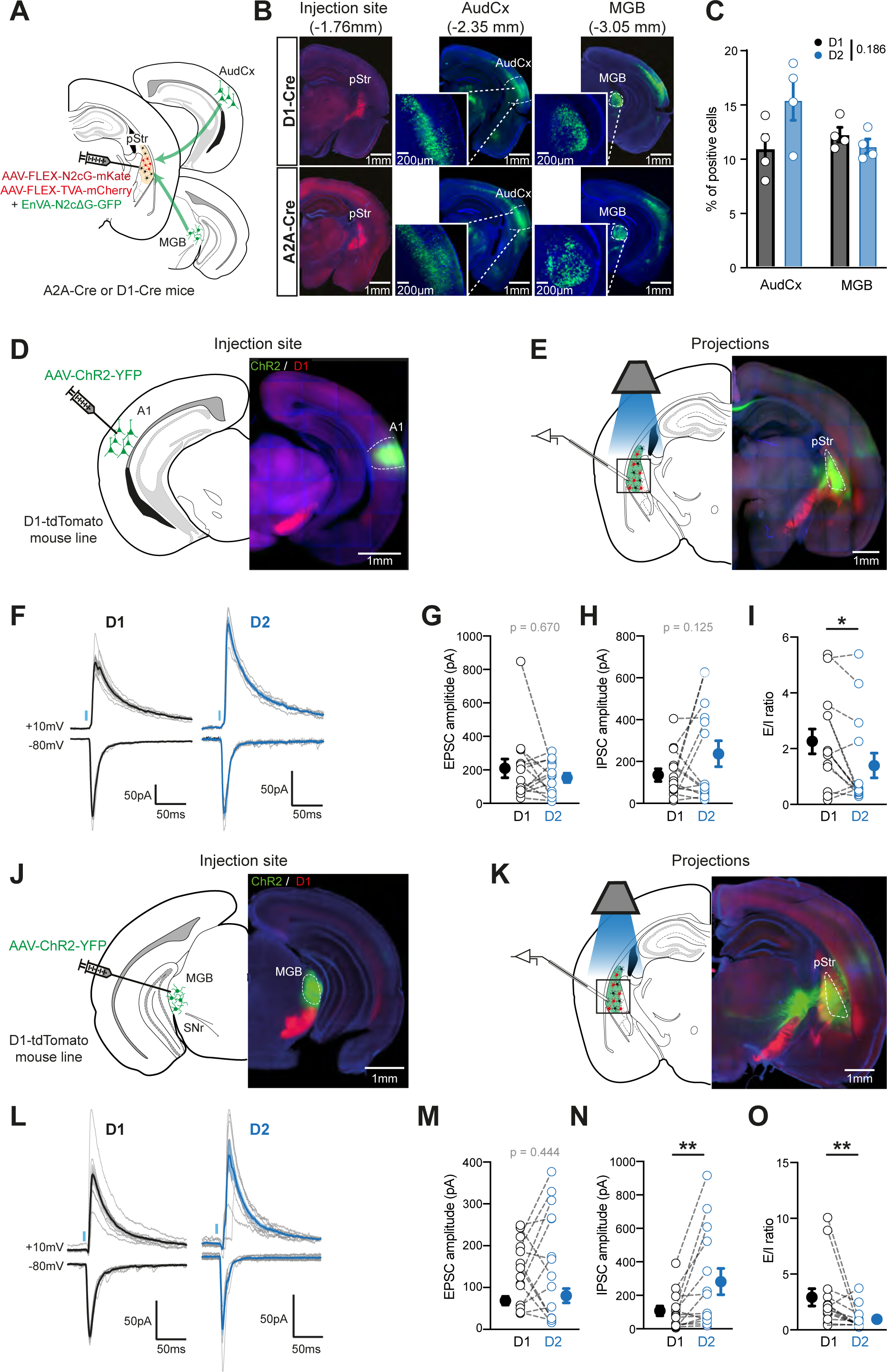
Cell-type specific connectivity from thalamus and cortex to the pStr. (A) Schematic illustrating rabies injection into the pStr of A2A-Cre or D1-Cre mice, labeling monosynaptic inputs from throughout the brain, including the auditory thalamus (MGB) and primary auditory cortex (A1). (B) Starter cells in the injection site were labeled with mCherry and mKate (red), while neurons monosynaptically connected were labeled with GFP throughout the brain. (C) The percentage of cells projecting from A1 and MGB onto D1 and D2 SPNs was similar. (N = 4 for both groups, Cell-type factor p = 0.287) (D) To enable optogenetic circuit mapping, A1 was injected with an anterograde AAV expressing ChR2. (E) In mice where A1 was injected with ChR2, D1 and D2 neurons were sequentially targeted for patch clamp in the region of pStr that showed the highest density of axons. (F) Examples of light-evoked excitatory postsynaptic currents (EPSCs) elicited at −80 mV (bottom) and inhibitory postsynaptic currents (IPSCs) recorded at +10 mV in D1 SPNs (left, black) and D2 SPNs (right, blue). Grey traces are individual trials. (G) A1 light-evoked EPSC amplitude was not significantly different between D1 and D2 SPNs (n = 14 pairs; p = 0.670). (H) A1 light-evoked IPSC amplitude was not significantly different between D1 and D2 SPNs (n = 14 pairs, p = 0.125). (I) The A1 light-evoked E/I ratio was larger in D1 than D2 SPNs (n = 14 pairs, p = 0.011). (J) same as (D) but with MGB injection. (K) same as (E) but with MGB projections. (L) same as (G) but with MGB injection. (M) MGB light-evoked EPSC amplitude was not significantly different between D1 and D2 SPNs (n = 14 pairs, p = 0.44) (N) MGB light-evoked IPSC amplitude was significantly larger in D2 compared to D1 and D2 SPNs (n = 14 pairs, p = 0.003). (O) The A1 light-evoked E/I ratio was larger in D1 than D2 SPNs n = 14 pairs, p = 0.007). All data are represented as mean ± SEM. * p < 0.05, ** p < 0.01.

Having established that there is no difference in the extent to which MGB and AudCx innervate D1 and D2 SPNs in the pStr, we next set out to test the relative strength of these connections. First, in D1-tdTomato or D2-Cre x Ai14 mice, we expressed Cre-independent ChR2 in A1 (Figure 2D-E). Weeks later, we obtained voltage-clamp whole-cell recordings in brain slices from D1 or D2 SPNs (identified by the presence or absence of tdTomato) under DIC optics. Recordings were made from 14 pairs of neighboring (within 80 µm) D1 and D2 SPNs patched sequentially (Figure S2E-F). Brief (8 ms) flashes of blue light resulted in excitatory postsynaptic currents (EPSCs) when measured at a holding potential of −80 mV and inhibitory postsynaptic currents (IPSCs) at +10 mV in both D1 and D2 SPNs (Figure 2F). IPSCs evoked by either A1 and MGB stimulation were the result of a feed forward inhibitory circuit, as they were both blocked by the glutamatergic antagonists CNQX and APV (Figure S2G and S2J). Comparing ChR2-evoked EPSCs in D1 and D2 SPNs revealed no significant difference in amplitude (Figure 2G; D1 = 209.4 ± 56 pA, D2 = 152.8 ± 27 pA, p = 0.67; Wilcoxon matched-paired test). IPSC amplitudes were also not statistically different between D1 and D2 SPNs (Figure 2H; D1 = 134.8 ± 30 pA, D2 = 235.7 ± 62 pA, p = 0.12; Student’s paired t-test). However, the ratio of the EPSC to IPSC amplitudes (E/I ratio) was statistically larger onto D1 SPNs compared to D2 SPNs (Figure 2I; D1 = 2.27 ± 0.44, D2 = 1.41 ± 0.45, p = 0.011; Wilcoxon matched-paired test).

In separate animals, we infected MGB neurons with ChR2, and the resulting EPSC and IPSC responses were measured in 14 pairs of D1 and D2 SPNs (Figure 2J-L). As with A1, we observed no difference in EPSC amplitudes (Figure 2M; D1 = 137.3 ± 19.9 pA, D2 = 161.0 ± 34.0 pA, p = 0.444; Student’s Paired t-test). However, MGB-evoked IPSCs were larger in D2 SPNs compared to D1 SPNs (Figure 2N; D1 = 104.6 ± 28.8 pA, D2 = 279.1 ± 78.5 pA, p = 0.003; Student’s Paired t-test), resulting in a significantly greater E/I ratio onto D1 SPNs (Figure 2O; D1 = 2.92 ± 0.78, D2 = 0.97 ± 0.26, p = 0.0067; Wilcoxon matched-paired test). We observed no difference in the probability of connection from either A1 or MGB onto D1 or D2 SPNs, and IPSCs were confirmed to be of polysynaptic origin (Figure S2G-L). Overall, our results reveal that, despite receiving similarly strong excitation from auditory afferents, D1 and D2 SPNs differ in the strength of feed forward inhibition, with D2 SPNs receiving more inhibition than D1 SPNs.

### Feed forward inhibition by PV neurons contributes to cell-type specific auditory responses in striatum

Given that D1 SPNs have a higher E/I ratio compared to D2 SPNs, it is possible that this difference contributed to the larger depolarizations of D1 SPNs to auditory stimuli we observed in vivo. To test this ex vivo, we sequentially patched D1 and D2 neurons but now in current-clamp mode to assess the overall impact of auditory afferent optogenetic stimulation on the membrane potential and/or spiking of D1 and D2 SPNs. In line with our in vivo findings, D1 SPNs were found to be less excitable than D2 SPNs (Figure S3). Despite this, ChR2-evoked depolarizations were larger in D1 SPNs than D2 SPNs (Figure 3A-B; D1 = 11.51 ± 2.4 mV, D2 = 6.42 ± 1.34 mV, n = 15 pairs, p = 0.01; Student’s Paired t-test), so much so that APs were only observed in D1 SPNs. We next blocked feed forward inhibition with gabazine, a GABA_A_ antagonist. This resulted in comparable A1-evoked responses between D1 and D2 SPNs (Figure 3C-D; D1 = 8.78 ± 1.91 mV, D2 = 7.96 ± 1.55 mV, n = 16 pairs, p = 0.607; Student’s Paired t-test). We obtained similar results when we sequentially patched D1 and D2 SPNs in brain slices in which MGB terminals were infected with ChR2. Here too, the size of the evoked response was larger in D1 SPNs (Figures 3E-F; D1 = 10.47 ± 1.53 mV, D2 = 7.165 1.11 mV, n = 15 pairs, p = 0.0068; Student’s Paired t-test), a difference that was also blocked by gabazine (Figures 3G-H; D1 = 13.87 ± 2.56 mV, D2 = 14.47 ± 2.51 mV, n = 15 pairs, p = 0.264, Student’s Paired t-test).

**Figure 3:**
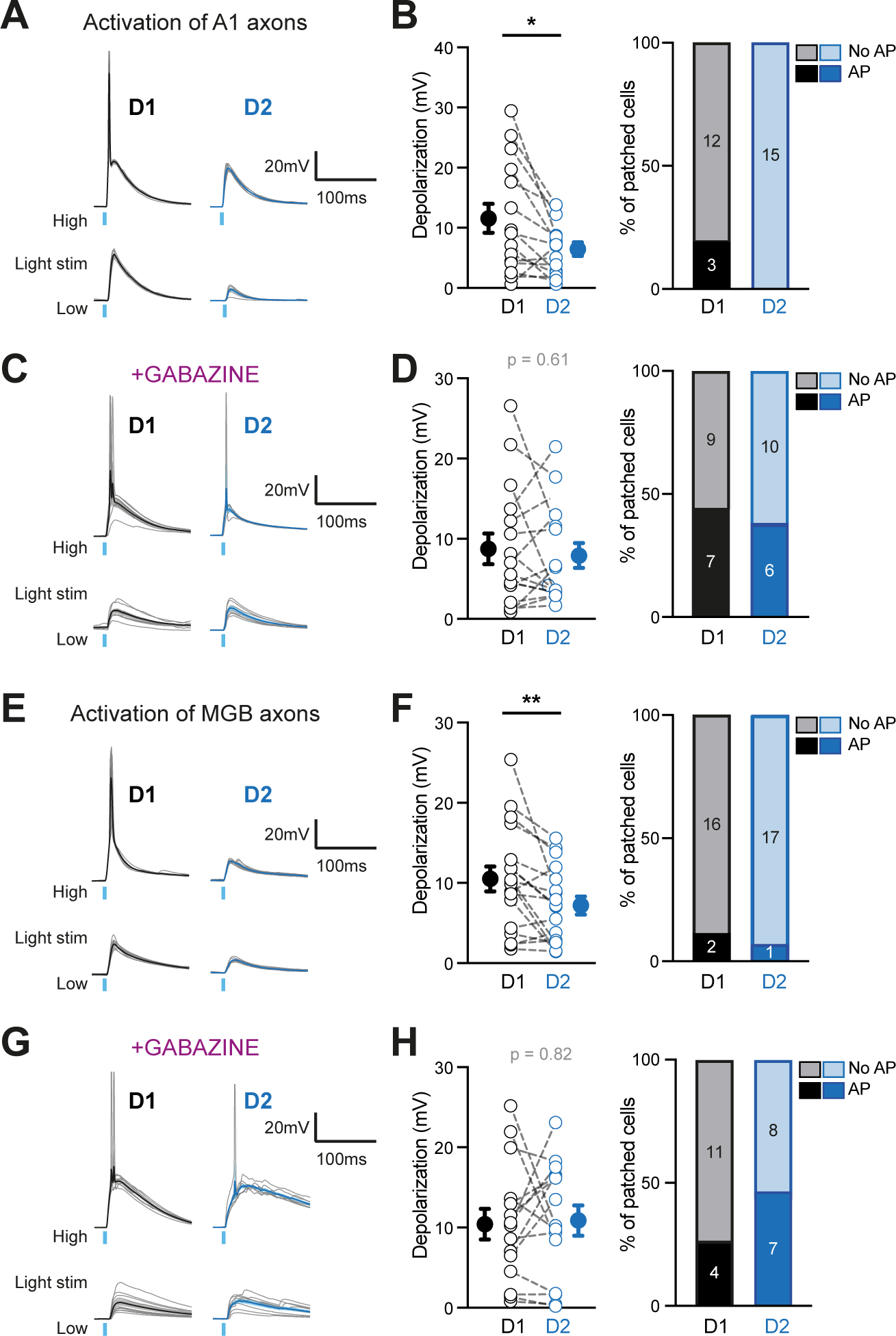
Feed forward inhibition from MGB and A1 contributes to cell type-specific evoked activity. (A) Example traces of A1 light-evoked PSPs with high (top) and low (bottom) light intensity in D1 (left, black) and D2 SPNs (right, blue). Example traces shown in grey. (B) Left: D1 SPN light-evoked PSP amplitude was significantly larger compared to D2 SPNs (n = 15 pairs; p = 0.010). Right: only D1 SPNs fired APs in response to light stimulation (n = 15 pairs, p = 0.068). (C) Same as (A) but with inhibitory transmission blocked with gabazine. (D) Left: With inhibitory transmission blocked, the A1-evoked PSP amplitude is not different between D1 and D2 SPNs. (n = 16 pairs, p = 0.607). Right: AP probability is also similar (n = 16 cells, p = 0.720). (E) Same as (A) but with MGB stimulation. (F) Same as (B) but with MGB stimulation. Left: n = 18 pairs, p = 0.0068. Right: AP probability in response to light was not significantly different between D1 and D2 SPNs (n = 18 pairs, p = 0.546). (G) Same as (C) but MGB stimulation (H) Same as D but with MGB stimulation (n = 15 pairs) Right: AP probability is also not different (n = 16 cells, p = 0.256). All data are represented as mean ± SEM. * p < 0.05, ** p < 0.01.

While several factors could underlie the difference in E/I ratio we observed between D1 and D2 SPNs, fast spiking interneurons (FSIs) are well known for exerting powerful and fast feed forward inhibition in the striatum^30,31^. We therefore wondered whether they differentially innervate D1 and D2s. To do so, we analyzed recordings in which one of the recorded tdTomato-negative neurons was an FSI, recognizable by its distinct firing properties and narrow spike width. Notably, these neurons exhibited a high rate of connectivity to both A1 and MGB, and were easily driven to fire APs upon auditory afferent stimulation, consistent with other studies^4,32^ (Figure S4A-C). Having established that FSIs in the pStr receive robust synaptic connections from MGB and A1, we next set out to test the strength the connections these neurons make onto D1 and D2 neurons. For this, because FSIs in the striatum are known to express pavalubumin (PV) we expressed ChR2 in PV-Cre mice^33^, and made ex vivo slices in which we sequentially patched D1 and D2 SPNs in voltage-clamp at +10 mV while briefly (8 ms), stimulating PV neurons with blue light (Figures 4A, S4D-F). These experiments revealed a larger PV neuron-evoked IPSCs onto D2 SPNs (Figure 4B-C; D1 = 90.3 ± 11.3 mV, D2 = 148.1 ± 22.4 mV, n = 21 pairs, p = 0.0251; Student’s Paired t-test). In addition, the paired pulse ratio (PPR) was lower in D2 SPNs, indicating a higher initial probability of release at these synapses (Figures 4D-E; 2^nd^, D1 = 0.859 ± 0.039; D2 = 0.741 ± 0.019, p = 0.052, 3^rd^, D1 = 0.731 ± 0.030; D2 = 0.577 ± 0.030; n = 7 pairs, p = 0.0035; cell-type factor p = 0.0056 Two-Way RM ANOVA).

**Figure 4:**
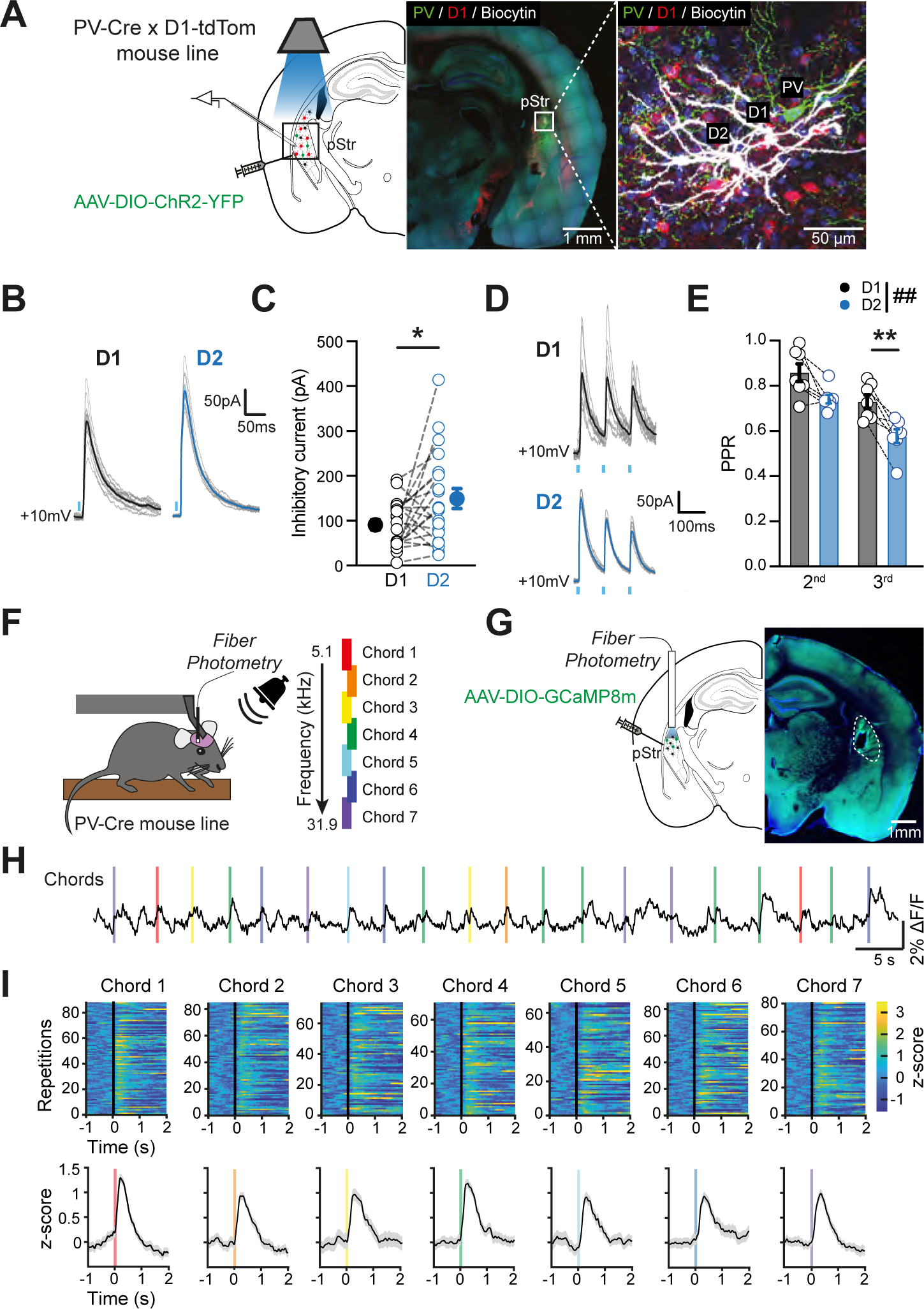
PV neurons in the pStr connect more strongly onto D2 SPNs. (A) Optogenetic circuit mapping PV+ neuron connectivity onto D1 and D2 SPNs in pStr was performed by injecting AAV-DIO-ChR2-YFP in PV-Cre x D1TdTom mice. D1 SPNs expressing tdTomato, and D2 SPNs were sequentially patched and filled with biocytin in the vicinity of a PV neuron expressing ChR2. (B) Example traces of PV light-evoked IPSCs in D1 (left, black) and D2 SPNs (right, blue), grey traces are individual trials. (C) PV light-evoked IPSC amplitudes were larger in D2 SPNs than in D1 SPNs (n = 21 pairs; p = 0.0251). (D) Example traces of PV light-evoked IPSCs in response to trains of stimuli (10Hz) in D1 SPNs (top, black) and D2 SPNs (bottom, blue), individual trials shown in grey. (E) The paired-pulse ratios showed a larger depression in D2 SPNs than in D1 SPNs (n = 7 pairs; Two-way RM ANOVA Cell-type factor: p = 0.0056). (F) Mice were head-restrained for fiber photometry recording of calcium activity of PV interneurons during sound presentation of the 7 discrete chord stimuli as presented in Fig.1. (G) AAV-DIO-GCaMP8m-YFP was injected into the pStr of PV-Cre mice, allowing for expression of ChR2 in PV+ neurons. (H) Example photometry trace showing fluorescence fluctuations representing activity of PV neurons in these mice. Vertical colored bars indicate time of auditory stimulus presentations. (I) Top: Plot showing z-scored Ca^2+^ signals in all trials records from all mice (n = 4), aligned to the stimulus onset (vertical black line). Bottom: Average z-score of Ca^2+^ activity in response to across all trials for each stimulus in these mice. All data are represented as mean ± SEM. ## p < 0.01 for Two-Way RM ANOVA ; * p < 0.05, ** p < 0.01

We wanted to confirm that PV interneurons are responsive to auditory stimuli in vivo. Since these neurons would only rarely be encountered using our whole-cell approach, we instead utilized PV-Cre mice to express the genetically encoded calcium indicator, GCaMP8m, specifically in these neurons. We then used fiber photometry to record the responses of PV cells to auditory stimuli (Figure 4F-G). These neurons showed strong stimulus-evoked responses to all stimuli we presented to the mouse (Figure 4H-I, n = 4 mice). Taken together, these results suggest that PV interneurons are readily recruited by auditory stimuli, make stronger connections onto D2 SPNs, and therefore likely contribute to the differential recruitment of direct and indirect pathways to auditory inputs.

## Discussion

In this study, by using the whole-cell recording technique in vivo, we measured the subthreshold voltage activations (i.e. synaptic inputs) evoked by auditory stimuli in the pStr in awake and anesthetized mice. These recordings revealed a bias of these responses onto D1 SPN neurons of the direct pathway. Optogenetic circuit mapping ex vivo revealed differences in feed forward inhibition onto D1 and D2 SPNs, with D1 SPNs showing a higher E/I ratio and receiving weaker inhibition from FSIs as compared to D2 SPNs. Our results extend our current knowledge of how auditory pathways recruit SPNs in the pStr and put forward a circuit mechanism by which these cell types can be differentially recruited by auditory stimuli to mediate behavior.

### Cell-type specific sensory responses in the striatum across modalities and behaviors

Given that the striatum receives input from many sensory cortical and subcortical areas, it is interesting to consider how our results compare those of other modalities. Recordings made from neurons in the dorsolateral striatum, the region of the striatum most heavily innervated by whisker barrel cortex, revealed that whisker-evoked synaptic responses were significantly larger in direct pathway neurons^27,34^. V_m_ responses to visual stimuli have been studied in SPNs^27^, however not in a cell-type specific manner, leaving it unclear if visual inputs follow the same trend. It is worth noting that sensory inputs of a single modality can innervate multiple areas of the striatum. Primary auditory cortex, for example, also shows substantial innervation of the dorsomedial striatum^2^, and it is unclear if these inputs exhibit patterns of innervation onto the direct and indirect pathway neurons like those we demonstrate here for the pStr.

Several studies have examined how output pathways of the striatum are activated in response to sensory stimuli that trigger behavior. Previous work from our group demonstrated stronger somatosensory-triggered subthreshold responses in D1 SPNs both before and after mice learned to perform a whisker detection task^5,6^. Imaging studies that have shown larger cue-evoked activity in the direct pathway in the DLS during an auditory cue-triggered motor task^35^, and preferential activation of the this pathway in the pStr upon contact with a threatening stimulus^16^. Therefore, it is possible that D1 SPNs are specialized to mediate sensory-driven actions, especially after sensorimotor learning^12,15,23^, although more experiments that systematically measure sensory inputs across different modalities to this pathway before and after learning are needed to validate this hypothesis. In addition, while D2 SPN activations were of lesser magnitude in our study, they are present and can result in AP firing^36^. What role these responses play in behavior is yet to be fully elucidated although some studies point to freezing and avoidance responses under certain conditions^15,16^.

### Feed forward inhibition as a mediator of cell-type specific sensory responses in the striatum

While fast spiking, PV, interneurons make up a small percentage of neurons in the striatum, they are known to be readily recruited by cortical inputs^4^ and active during periods when SPNs are under the influence of strong cortical and subcortical as well and dopaminergic inputs, as is the case for movement and learning ^37–39^. Our data demonstrate that in the pStr, PV interneurons receive strong synaptic connections from primary cortical and subcortical regions and are readily recruited by auditory stimuli. Because they make abundant connections onto nearby SPNs, PV interneurons then can powerfully regulate the responsiveness of these neurons to incoming inputs^30,40^. In the case of the pStr, a region thought to be important for mediating avoidance, it is possible that increased activation of D1 SPNs, mediated by the differential connectivity of PV interneurons onto SPN subtypes, allows for navigation away from novel, and potentially threatening, stimuli^15^.

An important outstanding question is the role that dopamine might play in shaping the responsiveness of D1 and D2 SPNs to auditory stimuli, both before and after learning. Dopamine projections to this region are particularly active during the presentation of novel auditory stimuli^12,13,41,42^ and in response to aversive outcomes. Given that dopamine is thought to positively regulate the responsiveness of D1 SPNs to synaptic inputs, it is possible that dopamine inputs contribute to the skewed activation of D1 SPNs to novel stimuli such as the ones we presented to mice in this study. However, dopamine inputs to this region have also been shown to play a role in reward-based learning where they contribute to the responsiveness of SPNs. Given that the activity of PV neurons is modulated by learning^43^ and connectivity onto SPNs is rapidly altered after dopamine depletion^44^, It is possible that PV connectivity onto these cell types is also modulated during learning. Future work may examine how these processes work in parallel to determine how the output pathways of the pStr are differentially recruited during both innate and learned behaviors.

## Acknowledgements

This work was supported by the National Institutes of Health (R01 NS126391), the Brain and Behavior Research Foundation Career Award for Medical Scientists (BBRF CAMS 1018390) and the Whitehall Foundation (Research Project Grant #2021-12-091) to T.S., the Leon Levy Foundation Fellowship to M.D, and the NYU Clinical and Translational Research TL1 to M.K. CVS-N2c rabies viruses were produced by the Center for Neuroanatomy with Neurotropic Viruses, supported by P40 OD010996. We thank Nicolas Tritsch and Richard Tsien for comments on the manuscript, Anders Nelson and Akira Fushiki for technical help with rabies tracing, Susan Brenner-Morton from the Columbia Zuckerman Antibody Core. Darcy Peterka and Luke Hammond and the Columbia Zuckerman Imaging Core for help with anatomical tracing analysis, and Rui Costa for guidance on this project.

## Author contributions

T.S. and M.D. contributed to study design and wrote the paper. M.D. and M.K. performed in vivo electrophysiology experiments. M.D. conducted ex vivo electrophysiology experiments, analyzed this data and performed the in vivo fiber photometry experiments. C.C. performed rabies injections, histology, immunohistochemistry and imaging of rabies injected mice and M.D. and C.F. performed analysis using the BrainJ pipeline. M.K created a database for all in vivo data and wrote the custom code to analyze this data.

## Methods

### Key resources table

| Reagent or Resource | Source | Identifier |
| --- | --- | --- |
| **Antibodies** |  |  |
| Streptavidin, Alexa Fluor 647 conjugate | ThermoFisher | S21374 |
| Mouse anti-Parvalbumin | Millipore | MAB1572 |
| Rabbit anti-dsRed | Zuckerman Institute Antibody Core, Columbia University | RRID: AB_3086780 |
| Donkey anti-Rabbit Alexa 594 | ThermoFisher | A-21207 |
| Donkey anti-Mouse Alexa 647 | ThermoFisher | A-32787 |
| **Virus** |  |  |
| AAV5-CaMKII-hChR2(H134R)-eYFP | Addgene | 26969-AAV5 |
| AAV2-Ef1a-DIO-hChR2(H134R)-eYFP | UNC Vector Core/Deisseroth | AV4378P |
| AAV2-CAG-FLEX-N2cG-mKate2.0 | Janelia Farm | - |
| AAV-EF1a-FLEX-TVA-mCherry | Stanford Gene Vector and Virus Core | GVVC-AAV-067 |
| EnVA-N2cΔG-GFP | Thomas Jefferson University | N/A |
| pGP-AAV1-syn-FLEX-jGCaMP8m-WPRE | Addgene | 162375-AAV1 |

| Chemicals, peptides, and recombinant proteins |  |  |
| --- | --- | --- |
| Biocytin | Biotium | 90055 |
| TTX | Hello Bio | HB1035 |
| 4-AP | Hello Bio | HB1073 |
| SR95531 | Hello Bio | HB0901 |
| Experimental models: Mouse lines |  |  |
| D2-Cre (ER44) | UCD | RRID:MMRRC_032108-UCD |
| Adora2a-Cre (KG139) | UCD | RRID:MMRRC_036158-UCD |
| D1-Cre (EY262) | UCD | RRID:MMRRC_030989-UCD |
| Drd1a-tdTomato | Jackson Labs | RRID:IMSR_JAX:016204 |
| Ai32(RCL-ChR2(H134R)/EYFP) | Jackson Labs | RRID:IMSR_JAX:024109 |
| Ai14 (RCL-tdT) | Jackson Labs | RRID:IMSR_JAX:007914 |
| Ai9 (RCL-tdT) | Jackson Labs | RRID:IMSR_JAX:007909 |
| PV-Cre | Jackson Labs | RRID:IMSR_JAX:017320 |
| Software and algorithms |  |  |
| ImageJ | https://imagej.net/ | RRID:SCR_003070 |
| MATLAB | MathWorks | RRID:SCR_001622 |
| IgorPro | Wavemetrics | RRID:SCR_000325 |
| pClamp 11 | MolecularDevices | RRID:SCR_011323 |
| GraphPad Prism 8 | http://www.graphpad.com/ | RRID:SCR_002798 |
| BrainJ | Botta et al. 2020 | N/A |
| **Other** |  |  |
| Custom code (Matlab) | This paper | Zenodo |
| Custom code (IgorPro) | This paper | Zenodo |
| Optopatcher | A-M Systems | 663849 |

### Resource availability

Further information and requests for resources and reagents should be directed to and will be fulfilled by the lead contact, Tanya Sippy).

No new materials were generated in this study.

Data and code availability: Data and custom code used in analysis is available on zenodo (DOI: 10.5281/zenodo.10641633).

### Animals

All experiments were carried out with 6-16 week-old male and female mice in accordance with protocols approved by the NYU Langone Health (NYULH) Institutional Animal Care and Use Committee (protocol #PROTO201900059). D1-Cre, A2A-Cre and D2-Cre bacterial artificial chromosome (BAC) transgenic mice were obtained from Gene Expression Nervous System Atlas (GENSAT; founder line EY262 for D1-Cre, KG139 for A2A-Cre and ER44 for D2-Cre), and purchased through the Mutant Mouse Regional Resource Centers (MMRRC). PV-Cre mice were purchased from Jackson Labs. For in vivo optopatching experiments, D2-Cre mice were crossed with lox-stop-lox (LSL) Channelrhodopsin reporter mice (Ai32, JAX: 012569) to obtain D2-Cre x LSL-ChR2 mice. For ex vivo slice experiments, Drd1a-tdTomato (JAX: 016204), D2-Cre and PV-Cre were crossed with Lox-Stop-Lox-tdTomato mice (Ai9, JAX: 007909 and Ai14, JAX:007914).

The mice were housed in a reverse light/dark cycle (light 11 pm to 11 am), at a temperature of 22 ± 2°C with food available ad libitum.

### In Vivo Electrophysiology

D2-Cre x Ai32 mice were implanted with a light-weight metal head post under isolflurane anesthesia. Seven to ten days later, mice underwent habituation to head fixation for 3 days in increasing time increments. On the day of the recordings, a small (less than 1mm diameter) craniotomy made under isoflurane anesthesia over the pStr (stereotaxic coordinates: 1.7mm posterior and 3.0mm lateral to bregma). For awake recordings, mice were allowed to recover from anesthesia for two to four hours, then transferred directly to the setup. For anesthetized recordings, mice were first injected with a mix of Ketamine/Xylazine (100 mg/kg; 10 mg/kg). Afterward, whole-cell patch clamp recordings were made as previously described^45^. 6-8 MΩ glass pipettes were filled with a solution containing (in mM): 135 K-methyl sulfonate, 5 KCl, 0.1 EGTA-KOH, 10 HEPES, 2 NaCl, 5 MgATP, 0.4 Na2GTP, 10 Na2-phosphocreatine, to which 2-4 mg/ml of biocytin was added. All patch clamp recordings in vivo were obtained in current-clamp mode without injection of any current, except during the characterization of intrinsic electrophysiological properties and the V_m_ was not corrected for liquid junction potentials. V_m_ signals were amplified using a Multiclamp 700B amplifier but digitized at 20 kHz using National Instruments acquisition boards (BNC 2110) and recorded in a MATLAB software (Wavesurfer, HHMI Janelia Research Campus).

At the start of each recording, a series of increasing current steps from −200 with increments of +25pA as injected into each neuron. The optopatcher was used for the online identification (A-M systems, WA USA). Light steps of 500 ms were delivered via a 470 nm LED (Thor labs) through an optic fiber (power at tip of the fiber was 1.5-3 mW) inserted into the patch-pipette while recording the responses in whole-cell configuration. Positive cells responded to light pulses with step-like depolarization, often exhibiting AP, while negative cells displayed no response or a small hyperpolarization. We proceeded with the recording if the neuron displayed both a stable resting V_m_ and overshooting action potentials. The series resistance of the recordings was between 25 to 50 MΩ. Our measurements of V_m_ in SPNs during sound presentation included between 4 to 25 trials for each chord with 3-5 s inter-trial intervals.

### Optogenetic Circuit Mapping In Ex Vivo Slices

Drd1a-tdTomato x PV-Cre mice were injected under isoflurane anesthesia with an adeno-associated virus (AAV) serotype 5 expressing double-floxed inverted reading frame humanized ChR2 (H134R) fused to YFP under control of CAMKII promoter (AAV5.CaMKII.hChR2(H134R).eYFP, virus made by Addgene with a titer of 2.3×10^10^ vg/mL). Prior to injection, a small ∼0.5 mm craniotomy was made over the area of the A1 (stereotactic coordinates: 2.30 mm posterior, 4.10 mm lateral to bregma, at a depth of 0.80 mm below the pia), MGB (stereotactic coordinates: 3.05 mm posterior, 1.85 mm lateral to bregma, at a depth of 3.08 mm below the pia), or the pStr (stereotactic coordinates: 1.58 mm posterior, 3.08 mm lateral to bregma, at a depth of 3.0 mm below the pia). A glass injection pipette was tip filled with the virus solution and lowered into the targeted brain area. 20-150 nl of the virus solution was slowly injected with a flow rate of 10 nl/sec. The micropipette was left in position for 5-10 minutes then slowly retracted to prevent backflow of the virus along the shaft.

2-3 weeks after viral injection, mice were deeply anaesthetized with isoflurane (5 % for 5-8 min) and perfused transcardially with 10 mL of ice-cold cutting solution containing (in mM): 110 Choline chloride, 2.5 KCl, 25 glucose, 25 NaHCO3, 1.25 NaH2PO4, 0.5 CaCl2, 7 MgCl2, 11.6 L-ascorbic acid and 3.1 sodium pyruvate. The brain was rapidly extracted and acute 300 μm coronal slices were cut in this solution with a vibratome (Leica VT1200S) then transferred to artificial cerebrospinal fluid (ACSF) containing the following (in mM): 125 NaCl, 2.5 KCl, 25 glucose, 25 NaHCO3, 1.25 NaH2PO4, 2 CaCl2, and 1 MgCl2, Slices were incubated in ACSF at 32°C for 20 minutes and then stored for at least 1 hour at room temperature. All solutions were constantly bubbled with 95% O2/5% CO2. For corticostriatal and thalamostriatal optogenetics, the cortex or thalamus was also sectioned and inspected to ensure that the ChR2 was targeted to A1 or MGB. If injections missed the target nucleus, slices were not utilized for recordings.

Targeted whole-cell recordings were made from neurons in the pStrr in a region that was densely innervated by A1 or MGB as visualized by expression of YFP axons. SPNs were first identified visually by differential interference contrast (DIC) imaging. The cellular identity of targeted neurons was confirmed through expression or lack of expression of transgenically-expressed reporters. D1 and D2 SPNs were sequentially patched, and cells were located at the same depth in the slice, less than 80 μm of each other (Figure S2D), and the recording order alternated between pairs.^46^ Current-clamp and voltage-clamp recordings were performed using borosilicate pipettes (3-5 MΩ) filled with potassium- or cesium-based intracellular solution containing the following (in mM): 135 K- or Cs-methyl sulfonate, 5 KCl, 0.1 EGTA-KOH, 10 HEPES, 2 NaCl, 5 MgATP, 0.4 Na2GTP, 10 Na2-phosphocreatine and 2-4 mg/ml of biocytin. Voltage or V_m_ signals were amplified using a Multiclamp 700B amplifier (Axon Instruments), digitized at 20 kHz by a Digidata 1550B AD/DA board (Molecular Devices) and acquired with the pClamp 10 software (Molecular Devices).

ChR2-expressing A1 and MGB axons were stimulated optically with 8 ms-long 470 nm light from a blue light-emitting diode (CoolLED pE-4000) through a 40 x 0.8 NA objective with a power range of 0.014 – 1.5 mW/mm^2^. For voltage-clamp experiments, responsive cells were defined by the presence of an EPSC at −80 mV in voltage-clamp configuration. Non-responsive cells were defined as not displaying a light elicited current at the maximum of the LED power. Pairs were only recorded when both cells were connected, and the same light was used for each pair. For characterization of light-evoked EPSCs and IPSCs, 6-10 trials were averaged.

For current-clamp experiments, cells were held at their resting membrane potential, responsive cells were defined as having a PSP in response to light stimuli and the same light intensity was used for each pair. AP probability was tested in each recorded cell with lights steps ranging from 0-100% power, and a cell was included as having APs if it fired at any of light intensity. Membrane resistances were determined by measuring the response to a voltage step command from −65 to −75 mV. Spike frequency-depolarization curves were generated by injecting series of 500 ms depolarizing current steps increasing by 25 pA.

For experiments mapping FSI (PV+) connectivity, SPNs were targeted for sequential patching as described above. Light-evoked IPSCs from PV neurons expressing ChR2 were elicited using 2 ms-long 470 nm light pulses through a 40 x 0.8 NA objective with a power range of 0.014 – 0.1 mW/mm^2^ and the same power level was used for each pair. For Paired-pulse Ratio (PPR) experiments, 10 Hz light trains at minimum light stimulation was used to elicit trains of stimuli.

To determine monosynaptic input, 1 µM tetrodotoxin (TTX, HelloBio) to prevent sodic AP generation and 100 µM 4-aminopyridine (4-AP, HelloBio) to block K+ voltage-dependent channels were added to ACSF. To determine the specificity of GABAergic currents at +10 mV, 10 µM of SR95531 (HelloBio) was added to the bath.

### Fiber Photometry

PV-Cre mice were injected in the pStr with 300 nl of AAV virus expressing double-floxed inverted reading frame GCaMP8m (pGP.AAV1.syn.FLEX.jGCaMP8m.WPRE) following the same procedure as describe above. A commercially available optic fiber (400 μm diameter; numerical aperture (NA) = 0.5 ; RWD Life Science Co.) was implanted above the injection site and cemented to the skull with a head bar. To record the GCaMP8m signal, excitation light was passed through a fiber-optic patch cord (Doric, 400 μm, 0.48 NA) coupled to a LEDs at 470 nm (Thorlabs, M470F3) connected to a fluorescence mini-cube (Doric, FMC5_E1(460-490)_F1(500-540)_E2(555-570)_F2(580-680)_S). Excitation light was delivered in continuous wave with light power measured at the tip of the fiber-optic patch cord was set to 20–40 μW. Emission light was collected through the same patch cord and fluorescence mini-cube connected to a photoreceiver (Newport, 2151; set to DC mode). Photometry signals were digitized at 20 kHz by a National Instruments acquisition board. 3 weeks after the injection, mice underwent habituation to head fixation as described above. On the day of recording, mice were placed on the rig and photometry signals were recorded in response to each cord. Between 15 and 22 trials were collected for each chord.

### Retrograde Tracing

For rabies-based transsynaptic tracing from striatal neurons, a 60 nl of a 1:1 mixture of AAV-FLEX-N2cG-mKate (9.83×10^10^ GC/mL) and AAV-EF1a-FLEX-TVA-mCherry (9.3×10^12^ vg/mL) were injected into the pStr at a depth of 2.95–3.05 mm, 1.60-1.68 mm posterior and 3.12 mm lateral to bregma. Two weeks later, mice were injected in the same location with 350 nl of pseudotyped, G-defiicient rabies virus (2.0×10^9^ ffu/mL).

### Histology

Mice were deeply anaesthetized with isoflurane (5 % for 5-8 min, inhaled) and perfused transcardially with 4 % paraformaldehyde (PFA) in 0.1 M PBS. Brains were removed and fixed in 4 % PFA for a maximum of 24 hours in the same solution, which was then replaced by a 0.1 M PBS solution. 50 µm thick coronal slices were for used for rabies experiments and PV staining, and 100 µm coronal slices were cut for in vivo recorded brains (vibratome, Leica VT1000S). To identify PV neurons, slice were stained with anti-Parvalbumin antibody (1:1000, Millipore) overnight at 4°C, and incubated with a secondary antibody coupled with Alexa 647 (1:1000, ThermoFisher). To amplify the rabies injection site, slices were stained for dsRed (rabbit anti-dsRed, 1:16000, Zuckerman Institute Antibody Core, Columbia University) and incubated with a secondary antibody coupled with Alexa 594 (1:1000, Invitrogen). The slices were subsequently mounted with DAPI mounting media (Southern Biotech). Images were obtained with a custom-built Nikon AZ100 multizoom microscope, equipped with a 4x 0.4NA objective with an automated slide loader at the Columbia University Zuckerman Institute’s Cellular Imaging platform.

For in vivo experiments, slices were cut at 100 µm incubated with Streptavidin coupled to Alexa 647 (1:2000, Invitrogen). Images were obtained with a slide scanner (Olympus VS2000) with a 20x objective to identify the location of recorded cells.

### Quantification and Statistical Analysis

#### In vivo recordings

All data analysis of in vivo recordings data was performed in MATLAB using custom written algorithms. Rheobase was reported as the minimum current injection required to elicit an AP in each cell, as determined from a protocol in which increasing current steps were elicited in steps of 50 pA to each cell. Input resistance (R_in_) was measured as the slope of a linear fit between injected hyperpolarizing current steps (from –200 pA to 0 pA in steps of 50 pA).

To assess the auditory stimulus-triggered response, V_m_ changes to the same chord were evaluated relative to a baseline V_m_ averaged 200 ms before the stimulus and averaged. A neuron was considered responsive if the baseline-subtracted V_m_ in the period 0-50 ms after the onset of the chord presentation was at least 2 mV and has a z score greater than or equal to 3.

#### Ex Vivo recordings

Data analysis was performed in IgorPro (Wavemetrics) with custom-written code. Mean traces were calculated by averaging over 6 to 10 single trials. Voltage traces and membrane potential traces were aligned to the onset of the ChR2 stimulus of the MGB or A1 axons, or PV+ neurons. Mean PSC or PSP amplitudes were calculated by taking the average peak and subtracting the pre-stimulus voltage or potential. Baselines were defined as 300 ms before the stimulus onset. PPR was calculated as the ratio of the peak amplitude of the averaged current response evoked by the second, third and fourth stimulation to the peak amplitude of the averaged current response evoked by the first stimulation.

#### Automated Anatomical Reconstruction

Image processing and analysis was performed using BrainJ, a collection of custom tools developed to facilitate automated whole-brain analysis of tissue sections, as previously described.^47,48^ Briefly, tissue sections were first arranged anterior-posterior and processed to remove external fluorescence and neighboring objects. Subsequently, sections are centered and oriented to facilitate 2D rigid body registration^49^ to make a 3-D brain volume. Cell bodies were detected using Ilastik^50^, a machine learning-based pixel classification approach on images that had background fluorescence subtracted using a rolling ball filter. Data was output from the BrainJ pipeline the form of CSV files containing measurements of cell body counts from each region in the Allen Brain Atlas Common Coordinate Framework. These measurements were organized so that analyses from all brain sub-regions could be performed in the Allen Institute Mouse Common Coordinate Framework. Custom written code in MATLAB was then used to calculate the relative ratio of cells in each brain region and compare between D1-Cre and A2A-Cre injected animals.

#### Fiber Photometry

Raw photometry signals were processed using an available MATLAB code. In brief, raw voltage signals were down-sampled to 50 Hz (above the Nyquist frequency to prevent aliasing), and the final photometry signal (output as a percentage) was obtained using the equation ΔF/F = (F – F0)/F0, in which F0 is baseline fluorescence as describe (Krok, et al. 2023). The latter was computed by interpolating the bottom percentile of fluorescence values measured in 30-second-long sliding windows (0% overlap) along the entire photometry trace. Sensory responses to auditory stimuli were first sorted according to chord number and then averaged for all trials across the 3 mice within chords to calculate response amplitude.

#### Statistical Analysis

Data are presented as mean ± SEM. Statistical analyses were performed using Prism 9 (GraphPad) and MATLAB custom code. The normality of data distribution was tested using D’Agostino & Pearson test. For unpaired datasets, two-tailed Student’s *T*-tests (for normally distributed datasets) or Mann–Whitney tests (for nonnormally distributed datasets) were employed. For paired datasets, two-tailed Student’s Paired T-test or Wilcoxon Signed-Rank test (for nonnormally distributed datasets) were employed. For multiple comparisons, we used Two-Way ANOVA followed by Sidak’s test. Values of *P* < 0.05 were considered statistically significant. *P* values are reported as follows: \**P* < 0.05; \*\**P* < 0.01; \*\*\**P* < 0.001; \*\*\*\**P* < 0.0001.

