## Supplementary files for "Cell-type specific auditory responses in the striatum are shaped by feed forward inhibition"

Druart et al.

### Supplementary Figures and Legends

#### Figure S1

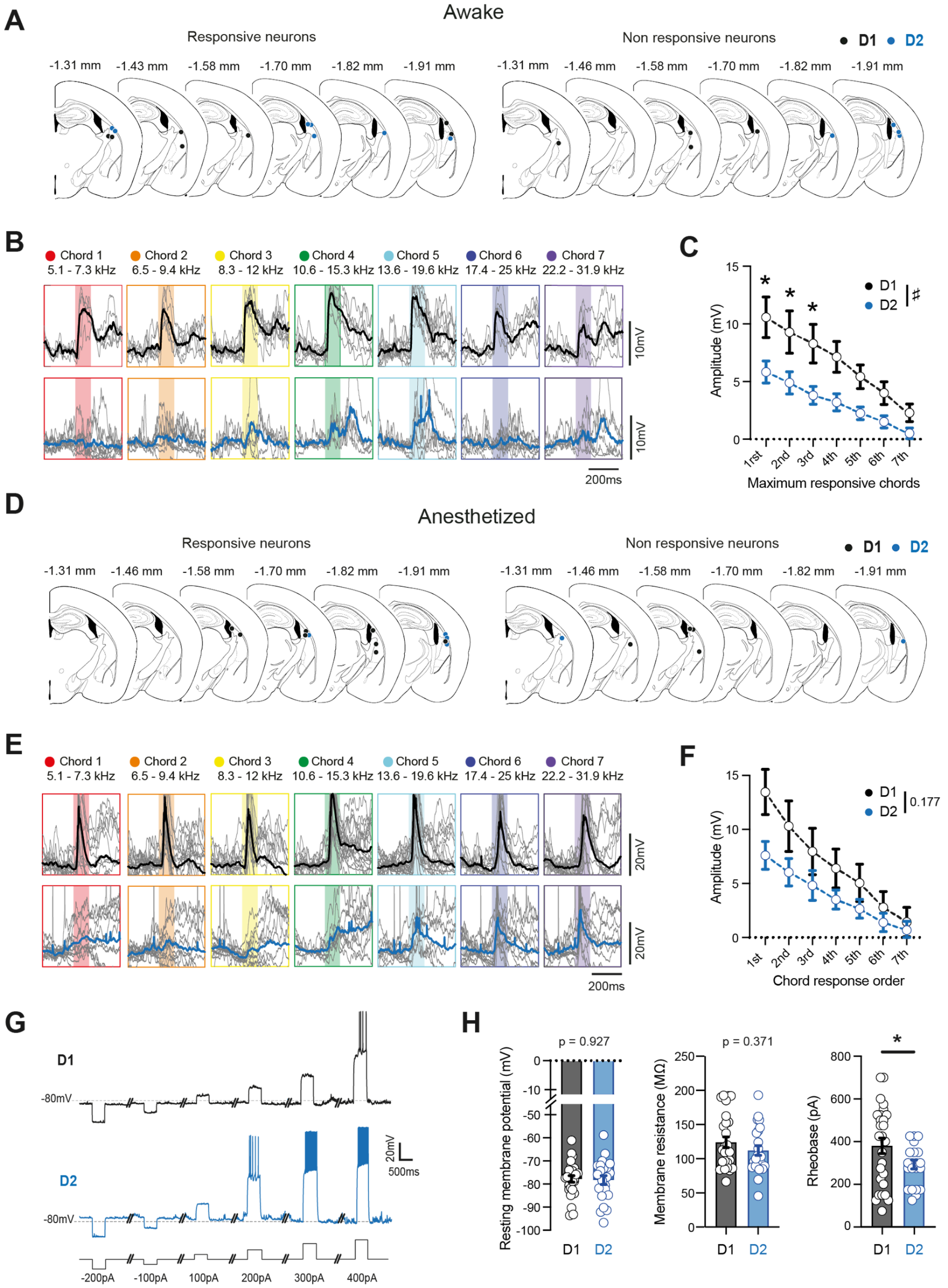

**Figure S1, related to Figure 1: Characterization of intrinsic properties and auditory responses of SPNs in vivo**

(**A**) Location of responsive (left) and nonresponsive (right) neurons from awake in vivo recordings. 13 D1 and 14 D2 SPNs representing 52% and 62% of the total data set were positively identified with post-hoc immunochemistry.

(**B**) Top: Baseline subtracted $V_m$ of the D1 SPN shown in Figure 1D for all trials in response to all 7 chords (black average, grey individual, chord 1, n = 5 trials; chord 2: n = 7 trials, chord 3: n = 8 trials, chord 4: n = 8 trials, chord 5: n = 10 trials, chord 6: n = 6 trials, chord 7: n = 6 trials). Bottom: Same for D2 SPN in Figure 1F, (blue average, grey individual, chord 1, n = 10 trials; chord 2: n = 8 trials, chord 3: n = 6 trials, chord 4: n = 5 trials, chord 5: n = 7 trials, chord 6: n = 9 trials, chord 7: n = 6 trials).

(**C**) Average amplitude of D1 SPNs (black) and D2 SPNs (blue) to the chord that yielded the maximum response ($1^{st}$) and the subsequent chords that yielded progressively smaller responses ($2^{nd}$ largest, $3^{rd}$, etc). D1 SPNs showed a larger response to more chords than D2 SPNs (D1: n = 12 cells; D2: n = 12 cells; # Two-Way RM ANOVA, cell-type factor : p = 0.0182; Sidak's post-hoc test, $1^{rst}$ : p = 0.021 ; $2^{nd}$ : p = 0.041 ; $3^{rd}$ : p = 0.035; $4^{th}$ : p = 0.089; $5^{th}$ : p = 0.281; $6^{th}$ : p = 0.577; $7^{th}$ : p = 0.876).

(**D**) Same as (E) but from recordings performed in anesthetized mice. 8 D1 and 7 D2 SPNs representing 53% and 63% of the total data set were positively identified with post-hoc immunochemistry.

(**E**) Top: Baseline subtracted $V_m$ of the D1 SPN shown in Figure 1M for all trials in response to all 7 chords (black average, grey individual, chord 1, n = 11 trials; chord 2: n = 20 trials, chord 3: n = 14 trials, chord 4: n = 22 trials, chord 5: n = 19 trials, chord 6: n = 13 trials, chord 7: n = 17 trials). Bottom: Same for D2 SPN in Figure 1O, (blue average, grey individual, chord 1, n = 12 trials; chord 2: n = 12 trials, chord 3: n = 12 trials, chord 4: n = 9 trials, chord 5: n = 9 trials, chord 6: n = 14 trials, chord 7: n = 11 trials).

(**F**) Same as (B) but from anesthetized recordings (D1: n = 10 cells; D2: n = 9 cells; Two-Way RM ANOVA, cell-type factor: p = 0.177).

(**G**) Example traces showing $V_m$ recording in a D1 SPN (black, top) and D2 SPN (blue, bottom) during step current injections from -200 pA to 400 pA.

(**H**) Left: D1 and D2 SPNs showed similar resting membrane potential (left, D1: -77.37 ± 1.44 mV, n = 27 cells; D2: -77.59 ± 2.01 mV, n = 21 cells, Student's t-test; p = 0.927). Middle: membrane resistance ($R_{in}$) did not show differences between D1 and D2 SPNs (D1: 123.3 ± 7.7 MΩ, n = 27 cells; D2: 113.3 ± 7.7 MΩ, n = 21, Student's t-test; p = 0.371). Right: D2 SPNs had a significantly lower rheobase than D1 SPNs (D1: 380.6 ± 36.0 pA n = 27 cells; D2: 285.7 ± 20.6 pA, n = 21 cells, Student's t-test; p = 0.039). Each open circle represents an individual cell.

All data are represented as mean ± SEM. # p < 0.05 for Two-Way RM ANOVA; * p < 0.05.

#### Figure S2

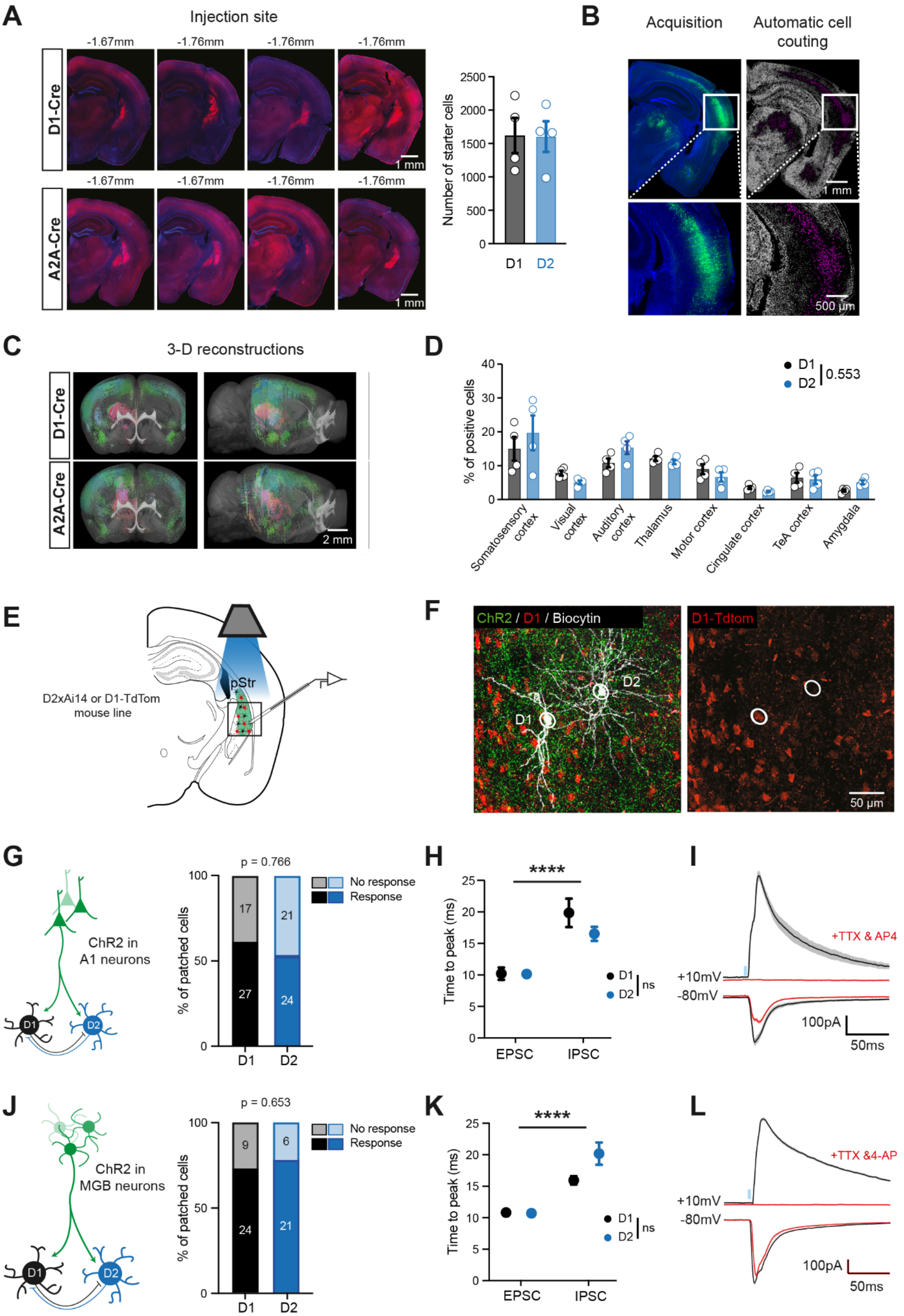
A
Injection site
-1.67mm
-1.76mm
-1.76mm
-1.76mm
D1-Cre
1 mm
-1.67mm
-1.67mm
-1.76mm
-1.76mm
A2A-Cre
1 mm
Number of starter cells
2500
2000
1500
1000
500
0
D1
D2
B
Acquisition
Automatic cell couting
1 mm
500 µm
C
3-D reconstructions
D1-Cre
A2A-Cre
2 mm
D
% of positive cells
40
30
20
10
0
D1
D2
0.553
Somatosensory cortex
Visual cortex
Auditory cortex
Thalamus
Motor cortex
Cingulate cortex
TeA cortex
Amygdala
E
D2xAi14 or D1-TdTom mouse line
pStr
F
ChR2 / D1 / Biocytin
D1
D2
D1-Tdtom
50 µm
G
ChR2 in A1 neurons
D1
D2
p = 0.766
% of patched cells
100
50
0
17
27
21
24
D1
D2
No response
Response
H
Time to peak (ms)
25
20
15
10
5
0
****
EPSC
IPSC
D1
D2
ns
I
+TTX & AP4
+10mV
-80mV
100pA
50ms
J
ChR2 in MGB neurons
D1
D2
p = 0.653
% of patched cells
100
50
0
9
24
6
21
D1
D2
No response
Response
K
Time to peak (ms)
25
20
15
10
5
0
****
EPSC
IPSC
D1
D2
ns
L
+TTX &4-AP
+10mV
-80mV
100pA
50ms

**Figure S2, related to Figure 2: Anatomy and physiology of A1 and MGB inputs to pStr D1 and D2 SPNs**

(**A**) Left: Rabies injection sites in the pStr of D1-Cre (top) and A2A-Cre (bottom) mice (n = 4 for both groups). Red cells are the starter cells. Right: Quantification of the number of starter cells per brain (D1-Cre in black and A2A-Cre in blue).

(**B**) Rabies-infected brains were sectioned, sorted and aligned to mouse brain coordinates with BrainJ software. Fluorescent images were then processed with Ilastik (see methods) to achieve automated cell counting (right, purple cells).

(**C**) 3-D reconstruction of a D1-Cre brain (top) and A2A-Cre brain (bottom) with all the starter cells (red) and projecting cells (green).

(**D**) Quantifications of the percentage of projecting cells from different brain regions in D1-Cre and A2A-Cre.

(**E**) Schematic for optogenetic circuit mapping experiments. D1 or D2 SPNs were identified based on their expression of tdTomato. Blue light through the objective was illuminated for brief periods (8 ms) to elicit ChR2-evoked responses from either MGB or A1 inputs.

(**F**) Histology from an experiment in which A1 axons were infected with ChR2-YFP and a D1 and D2 SPN were recorded in tandem near one another.

(**G**) Left: Simplified schematic of A1 to pStr connectivity. Right: percent of A1-stimulation responsive D1 and D2 SPN cells recorded in the pStr (Chi-square: $p = 0.766$).

**(H**) A1-evoked time to peak (TTP) of EPSCs was significantly shorter than time to peak of IPSCs in both D1 SPNs and D2 SPNs. (Two-Way ANOVA, $TTP_{EPSC}$ vs $TTP_{IPSC}$: **** $p < 0.0001$).

(**I**) Light-evoked EPSC (bottom) and IPSC (top) in the presence (red traces) and absence (black) of TTX (1 µM) and 4-AP (100 µM), showing that IPSCs are di-synaptic.

(**J**) Same as (E) but from MGB (Chi-square: $p = 0.653$).

(**K**) Same as (F) but from MGB.

(**L**) Same as (G) but from MGB.

All data are represented as mean ± SEM. **** $p < 0.0001$.

#### Figure S3

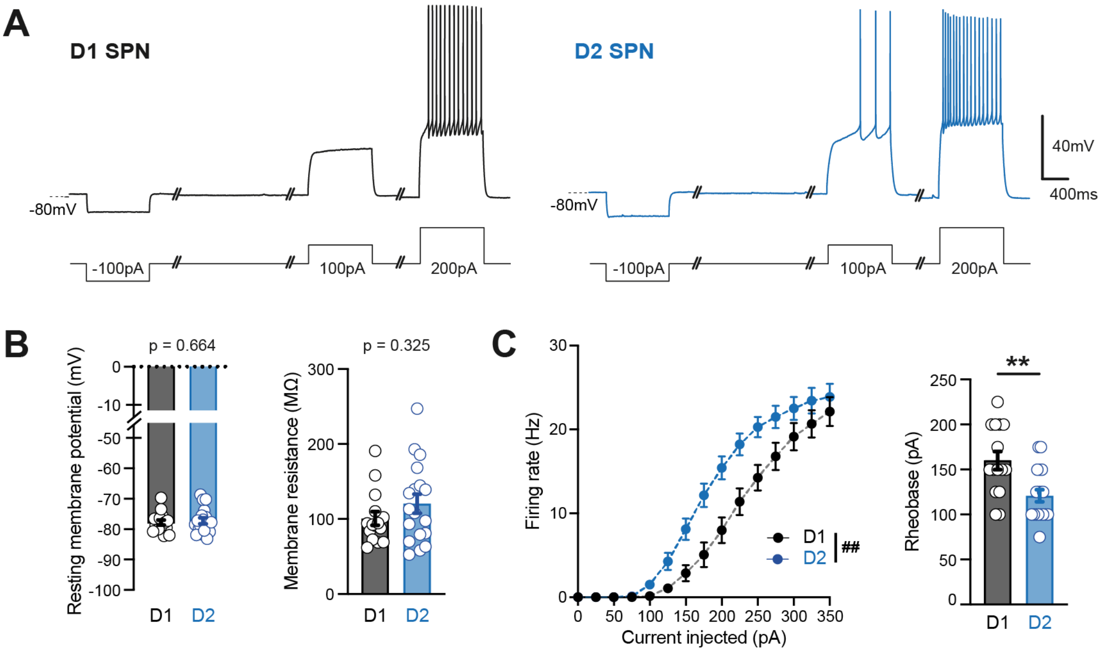


**Figure S3, related to Figures 2 and 3: Intrinsic properties of SPNs ex vivo**

(**A**) Example traces showing the $V_m$ in a D1 SPN (black, left) and D2 SPN (blue, right) during step current injections from -100 pA to 200 pA.

(**B**) Left: The resting $V_m$ in D1 and D2 SPNs was not significantly different (left, D1: n = 15 cells; D2: n = 18 cells; Student's t-test: p = 0.664) Right: The input resistance ($R_{in}$) was not significantly different between D1 and D2 SPNs (D1: n = 15 cells; D2: n = 18 cells; Student's t-test: p = 0.325).

(**C**) Left: F/I curves for D1 (black) and D2 SPNs (blue) showing that D2 SPNs are more excitable (D1: n = 15 cells; D2: n = 18 cells; ## Two-Way RM ANOVA, cell-type factor: p = 0.007). Right: D2 SPNs exhibit a significantly smaller rheobase than D1 SPNs (Student's t-test: p = 0.002).

All data are represented as mean ± SEM. ## p < 0.01 for Two-Way RM ANOVA; ** p < 0.01.

#### Figure S4

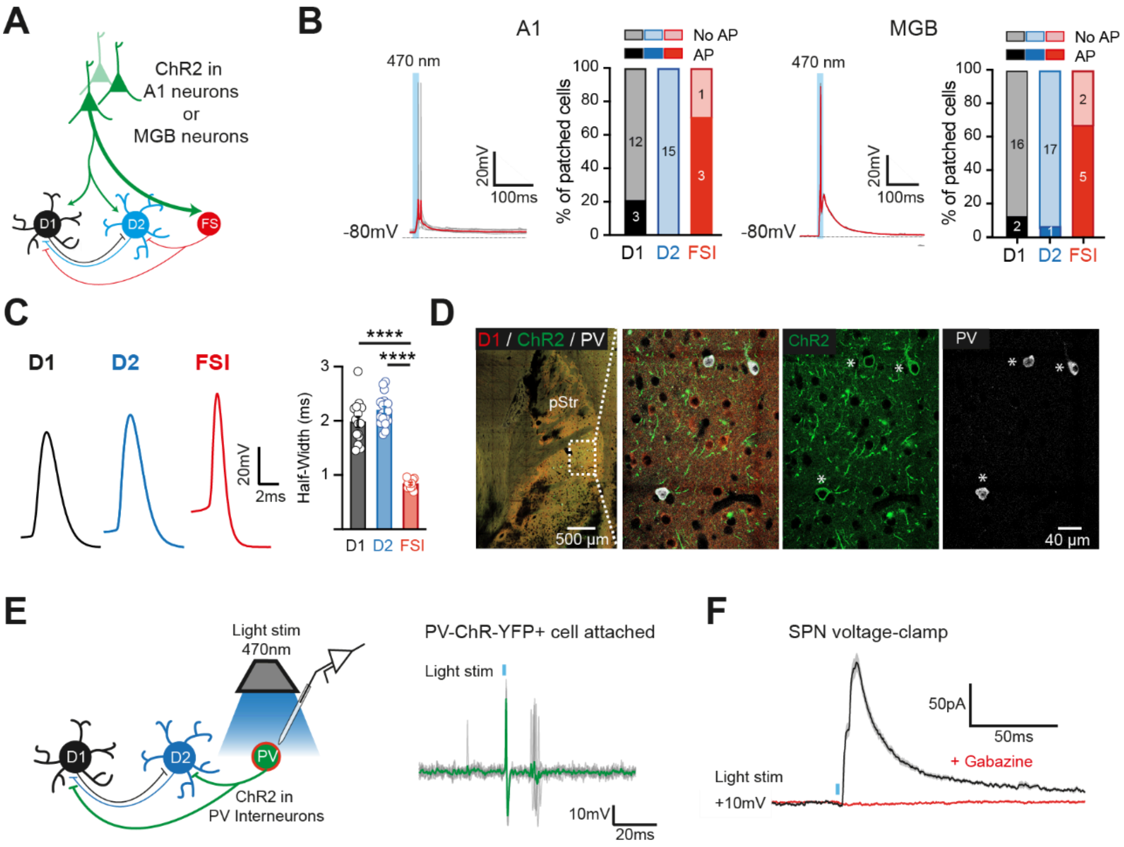


**Figure S4, related to Figure 4: Functional examination of PV interneurons in the pStr**

(**A**) Simplified schematic showing A1 and MGB connectivity into the pStr.

**(B)** Putative FSIs showed large and reliable PSPs in response to A1 (left, n =4 cells) or MGB (right, n = 7 cells) light evoked stimulation. Putative FSIs also more readily fired APs, compared to D1 or D2 SPNs.

**(C)** Putative FSIs were readily identifiable due to their narrow AP ½ width. This was significantly narrower than the ½ width of APs in D1 SPNs or D2 SPNs (#### One-Way ANOVA, $p < 0.0001$).

**(D)** In PV-Cre mice infected with AAV.DIO.hChR2-eYFP in the pStr, post-hoc antibody staining for PV neurons confirmed specificity of the ChR2-YFP expression in these neurons.

**(E)** Left, schematic showing experimental design in which ChR2-YFP PV neuron was targeted for cell-attached recording. Right: ChR2-expressing PV neuron in cell-attached configuration showing AP firing at light onset. Grey traces are individual trials and green is the average.

(**F**) Light-evoked IPSCs in a D1 SPN patched in a PV-Cre animal infected with AAV.DIO.hChR2-eYFP were blocked by the addition of Gabazine (10 µM), indicating the light-evoked response was mediated by GABAergic transmission.

All data are represented as mean ± SEM. **** $p < 0.01$ for One-Way RM ANOVA.
